# An unprecedented codon reassignment and two new genetic code variants in ciliates

**DOI:** 10.64898/2026.11.04.764322

**Authors:** Eliška Maturová, Vojtěch Žárský, Marek Eliáš

## Abstract

Ciliates are well-known for frequent nuclear genetic code changes. However, the exploration of ciliate nuclear genomes is an unfinished enterprise, and particular genetic code variants have been previously attributed to some ciliate lineages without solid evidence. We focused on two such groups, the class Litostomatea and deeply-branching lineages of the class Spirotrichea, and by analysing their genomes and transcriptomes we found that some of them employ genetic code variants differing from those suggested before. Particularly notable is the spirotrichean genus *Licnophora*, which we found to exhibit an unprecedented combination of codon reassignments, i.e. a new genetic code variant. In *Licnophora* spp., each of the three standard stop codons has a different altered meaning: UAA encodes glutamate, UAG glutamine, and UGA exhibits homonymy, encoding arginine when present in-frame and terminating translation at the coding sequence ends. Contrary to some previous claims, our analyses demonstrated that litostomateans generally employ the standard genetic code, with a single identified exception, the species *Balantidium ctenopharyngodoni*, which exhibits a codon reassignment never reported before: UGA encoding leucine. Whereas tRNAs decoding the reassigned UAA and UAG codons in *Licnophora* and UGA in *B. ctenopharyngodoni* were readily identified in the respective genomic data, the mechanism of UGA decoding as arginine in *Licnophora* and certain other spirotricheans remains elusive. Our results paint a more complex picture of genetic code evolution in Spirotrichea than assumed before and necessitate an update of the current list of genetic code variants to allow for an accurate prediction of protein sequences in ciliates.

**Significance Statement:** Accurate genome annotation and protein prediction critically depends on the genetic code of the focal organism to be correctly identified. Here, we demonstrate that two previously studied eukaryotes from the phylum Ciliophora in fact employ different genetic codes than assumed previously. Furthermore, both cases represent genetic code variants not yet reported from any organism. Namely, the litostomatean *Balantidium ctenopharyngodoni* is the first organism shown to use the UGA codon to encode leucine, and the deep-branching spirotrichean genus *Licnophora* has a unique stop codon reassignment setup, employing UAA, UAG and UGA to encode glutamate, glutamine and arginine, respectively. Our study thus further underscores the malleability of the genetic code and stresses the importance of explicit genetic code testing in genomic studies.

## Introduction

The outcome of translation of particular codons, i.e. the specific amino acid residue inserted in the nascent polypeptide chain or translation termination, is highly conserved across organisms and their translation systems. This correspondence is embodied in the so-called standard or universal genetic code. However, as early as in 1979 it was recognized that exceptions to this epitome of molecular biology exist, first discovered in the human mitochondrion (Barrell et al. 1979) and accumulating ever since then. A recent comprehensive review of genetic code variation attempted to provide an exhaustive list of known codon reassignments, i.e. changed meanings of particular codons, as well as of genetic code variants, i.e. specific combinations of reassigned codons employed by the given translation system (Lukeš et al. 2025). According to this census, over 50 different departures from the standard codon meanings, and roughly the same number of non-standard genetic code variants, have been documented in the literature.

Additions to the list of known genetic code changes come mainly from unanticipated encounters during continued genomic and transcriptomic exploration of the organismal diversity. However, changed codon meaning is not always easy to spot in the newly generated sequence data, or *a priori* assumptions about the codon meaning in the organism under scrutiny are frequently not properly tested in the original reports on the respective genomic or transcriptomic datasets. A common practice is to arbitrarily select a predefined genetic code variant (also called a translation table) from the list maintained by the National Center for Biotechnology Information (NCBI; https://www.ncbi.nlm.nih.gov/Taxonomy/Utils/wprintgc.cgi) and implemented in many programs in use, although this list is vastly incomplete and even includes translation tables misattributed to particular taxa or not supported by actual data at all (Lukeš et al. 2025). Indeed, numerous cases of the deployment of incorrect translation tables in published studies and sequence databases, effectively resulting in artificial amino acid sequences of the proteins encoded by the given genome, have been uncovered by rigorous reanalyses. Examples come from reassessments of the codon meaning in mitochondrial and plastid genomes of various protist groups that pointed to previously missed codon reassignments (Žihala and Eliáš 2019; Žihala et al. 2020; Barcytė et al. 2025), others were uncovered by systematic analyses of prokaryotic genomes, which detected the occurrence of previously unknown reassignments of sense codons in certain lineages of both Bacteria and Archaea (Shulgina and Eddy 2021; Melnykov 2026).

A systematic analysis of eukaryotic and organellar sequence data is so far lacking, and the space of available sequence data is expanding rapidly as well, so it is likely that its further exploration will deliver not only additional (independently evolved) instances of the known codon reassignments, but also unprecedented changes of the codon meaning or novel combinations of codon reassignments. For example, a recent study focusing on organellar genomes of a single algal group (Pedinophyceae) provided evidence for four previously unreported specific combinations of codon reassignments, necessitating the expansion of the defined translation table variants by four additional items to allow for correct conceptual translation of organellar coding sequences of the organisms concerned (Barcytė et al. 2025).

Codon reassignments and non-standard genetic codes are distributed very unevenly (Lukeš et al. 2025). They are rare in Bacteria (=Eubacteria), extremely rare in Archaea, sporadically present in plastids, yet very common and diverse in mitochondria. Regarding eukaryotic nuclear genomes interpreted by the cytoplasmic translation system, codon reassignments are primarily found in a few species-poor protist lineages representing terminal branches of the eukaryote tree of life (e.g., Záhonová et al. 2016; Pánek et al. 2017; Bachvaroff 2019; Pánek et al. 2025), along with three broader groups exhibiting an unusual evolutionary trajectory when it comes to the genetic code. In these three groups, namely Saccharomycotina (yeasts), ulvophyte algae, and ciliates, the genetic code seems to be inherently malleable, resulting in multiple parallel changes in each group, with varying outcomes (specific codon reassignments) evolved in different sublineages that are intermixed with sublineages exhibiting (primitively or potentially due to reversion) the standard genetic code. In yeasts, the changes affect the single codon CUG, normally coding for leucine but in different yeast lineages reassigned to serine or alanine (Mühlhausen et al. 2016; Krassowski et al. 2015), while in one species it is stochastically decoded as leucine or serine (Mühlhausen et al. 2018), exhibiting thus the so-called polysemy (see Lukeš et al. 2025). In ulvophytes, two different lineages having both UAG and UAA (normally termination codons) reassigned to encode glutamine are intermingled with a lineage exhibiting the standard genetic code and a lineage having UAG (but not UAA) reassigned to serine (Cocquyt et al. 2010; Li et al. 2021).

The most complex history of genetic code evolution among all eukaryotic groups is found in ciliates. Since the reports on the UAG and UAA codons encoding glutamine in nuclear genes of certain ciliates forty years ago (Caron and Meyer 1985; Hanyu et al. 1986), the number of different non-standard genetic code variants in ciliates has grown to more than ten, resulting from different combinations of twelve codon reassignments (Lukeš et al. 2025). Unlike the yeast case but analogously to codon reassignments in nuclear genetic codes of all other eukaryotes, the codon meaning changes in ciliates are (so far) restricted to the codons that normally serve as translation termination signals, i.e. UAG, UAA, and UGA. Hence, all the reassignments in ciliates are of the stop-to-sense category. However, the situation is complicated by the fact that some ciliate lineages exhibit an “incomplete” reassignment of some or all of these codons, whereby the interpretation of the codons by the translation apparatus depends on their position in coding sequences. When found within the coding sequence (CDS), these codons are translated as an amino acid, whereas their occurrences at the genuine 3’ end of the CDS are interpreted as a translation termination signal. This situation, recently categorized as a case of codon homonymy (sequence context-dependent alternative meaning of the same codon; Lukeš et al. 2025), was initially detected in a few ciliates and studied in depth in *Condylostoma magnum* (Swart et al. 2016; Heaphy et al. 2016), but seems to be much more widespread across the ciliate phylogeny (Seah et al. 2022; McGowan et al. 2023; Chen et al. 2023) and is also known to occur in a few eukaryotes outside ciliates (Záhonová et al. 2016; Bachvaroff 2019).

Some of the genetic code changes in ciliates have been reported only recently. Members of the class Oligohymenophorea are known to decode both UAG and UAA as the same amino acid (glutamine, except for the subclass Peritrichia decoding these codons as glutamate), but McGowan et al. (2023) discovered an uncultivated member of this class (Oligohymenophorea sp. PL0344) that decodes these codons as two different amino acids, UAG as glutamine and UAA as lysine. Furthermore, the latter reassignment had not been known from any translation system investigated before. Chen et al. (2023) reported another novel codon reassignment combination from *Peritromus kahli*, which decodes UAG and UAA as glutamine and UGA as cysteine, with UAA being homonymous (serving also as a termination codon). The same authors also proposed that some members of the class Spirotrichea exhibit dual (i.e., homonymous) decoding of UGA, with its in-frame instances translated as arginine (along with “fully” reassigned UAG and UAA translated as glutamine). Further recent additions to the list of ciliate genetic codes came with an investigation of nuclear genomes of representatives of the class Phyllopharyngea. Whereas the standard genetic code predominates in this group, two different sublineages were found to have reassigned the UAG codon to an amino acid (while retaining UAA as a stop codon): to glutamine in one sublineage and to leucine in the other (McGowan et al. 2024). The extreme propensity of ciliates to employ termination codons for amino acid encoding is indeed striking and points to a kind of instability in their translation system. It was proposed that the apparent evolutionary instability of the genetic code in ciliates reflect an (unknown) ancestral modification of their translation system that enables them to cope with “ambiguous” (or homonymous) meaning of termination codons before they get “fully” reassigned (Swart et al. 2016).

Given all these considerations, additional, so far missed codon reassignments and genetic code variants are likely to be discovered in ciliates that have not been (properly) investigated in this regard. Here we provide evidence that this is indeed the case, bringing to light two new genetic code variants including a codon reassignment not seen before.

## Results and discussion

### The unusual spirotrichean genus Licnophora exhibits a new genetic code variant

Two candidates to be checked for the specific status of the genetic code are *Licnophora macfarlandi* and *Phacodinium metchnikoffi*. To resolve their controversial phylogenetic position among ciliates, Irwin et al. (2021) generated transcriptome data from both species and employed them in a phylogenomic analysis, which resolved *L. macfarlandi* and *P. metchnikoffi* as two deepest branches in the ciliate class Spirotrichea (*L. macfarlandi* branching as a sister lineage to all other spirotrichs including *P. metchnikoffi*). In the description of their procedure the authors mentioned that the genetic codes selected for conceptual translation of the assembled transcriptome sequences were based on information provided in the NCBI Taxonomy database. As of June 2026, the NCBI database assigns the translation table 6, i.e. the widespread ciliate genetic code variant with UAG and UAA encoding glutamine, to *L. macfarlandi*, and the translation table 10, i.e. the “euplotid nuclear code”, to *P. metchnikoffi*. The basis of this assignment is not clear, as to our knowledge no explicit analysis of the codon meaning in these two ciliates has been published. The occurrence of the translation table 6 in *L. macfarlandi* would not be particularly surprising. However, assuming the “euplotid nuclear code”, which differs from the standard genetic code by UGA reassigned to cysteine and UAG and UAA serving homonymously as frame-shifting and translation termination signals (Lobanov et al. 2017; Chen et al. 2023), conflicts with a previously published claim (albeit not directly documented by specific data) that the genus *Phacodinium* uses UAG, UAA, and UGA as stop codons (Jiang et al. 2024). Furthermore, the spirotrichean family Euplotidae, in which the translation table 10 is known to occur, is not directly related to *P. metchnikoffi* (Irwin et al. 2021; Jiang et al. 2024).

To clarify the identity of the genetic code in both species, we assessed the most likely codon meaning in the two ciliates by analyzing their transcriptome assemblies with the dedicated tool Codetta (Shulgina and Eddy 2023); the results are presented in Table S1. Notably, we did not find any signal for UGA serving as a cysteine codon in *P. metchnikoffi*, and manual investigation of a sample of transcripts confirmed that UGA is used as a conventional termination codon, although rarely (Fig. S1). No indication for any other codon reassignment in *P. metchnikoffi* was observed either. Hence, consistent with the statement by Jiang et al. (2024), we conclude that this ciliate utilizes the standard nuclear genetic code instead of the translation table 10 indicated in the NCBI database and hence presumably employed by Irwin et al. (2021) in their analyses. Furthermore, the retention of the standard genetic code seems to be the rule for a broader *Phacodonium* lineage, as revealed by our analysis of multiple genome assemblies from the genome data available in the Protist 10 000 Genomes Project (P10K) database (Gao et al. 2024) assigned to this genus (Table S1).

A different picture was obtained for *L. macfarlandi*. Here the analysis with Codetta indicated a strong signal for UAG encoding glutamine, which would be consistent with the ciliate sticking to the translation table 6 as assumed by the NCBI database. However, the signal for UAA was unexpected yet very strong: this codon clearly encodes glutamate rather than glutamine in *L. macfarlandi*. Direct inspection of the UAG and UAA positions in a sample of coding sequences support the reassigned meaning of both codons as suggested by Codetta (Fig. 1A). An analogous variant, i.e. with UAG and UAA reassigned to two different amino acids, has been encountered once, in the uncultured Oligohymenophorea sp. PL0344, but as already stated above, in this ciliate the configuration of the codon meaning differs from the one found in *L. macfarlandi*, as UAA encodes lysine rather than glutamate; the genetic code alteration in *L. macfarlandi* is thus unique. To further illuminate its nature and origin, we used available genome sequencing reads from a related species *Liconophora lyngbycola* and obtained a partial genome assembly. Its analysis showed the same specific reassignment of the UAG and UAA codons as in *L. macfarlandi* (Fig. 1A; Table S1). Furthermore, in the genome assembly we identified genes for tRNAs carrying anticodons cognate to these two codons (Fig. 1B). Consistent with the inferred meaning of the UAG codon, the cognate tRNA_CUA_ most strongly matched the tRNA^Gln^ model when assessed by tRNAscan-SE 2.0 (Table S2), and our phylogenetic analysis showed it to belong to a cluster combining genes for standard tRNA^Gln^ species with tRNA^Sup^ genes from other spirotricheans (Fig. S2). In the case of the *L. lyngbycola* tRNA cognate to UAA, the prediction of the tRNA isotype was less conclusive, the expected tRNA^Glu^ model being only the fourth best (after Asp, His, and Thr), albeit with a score not being particularly worse that those of the three better-scoring isotypes (Table S2). Furthermore, the *L. lyngbycola* tRNA_UUA_ did not branch closely to any standard tRNA in our phylogeny (Fig. S2), making its origin unclear. Nevertheless, the signal for UAA encoding glutamate in *Licnophora*, as inferred from the occurrence of the codon at conserved positions by Codetta, is overwhelmingly strong when compared to the signal for any of the three amino acids slightly more preferred by the tRNA_UUA_ evaluation by tRNAscan-SE (Table S3).

**Fig. 1.**
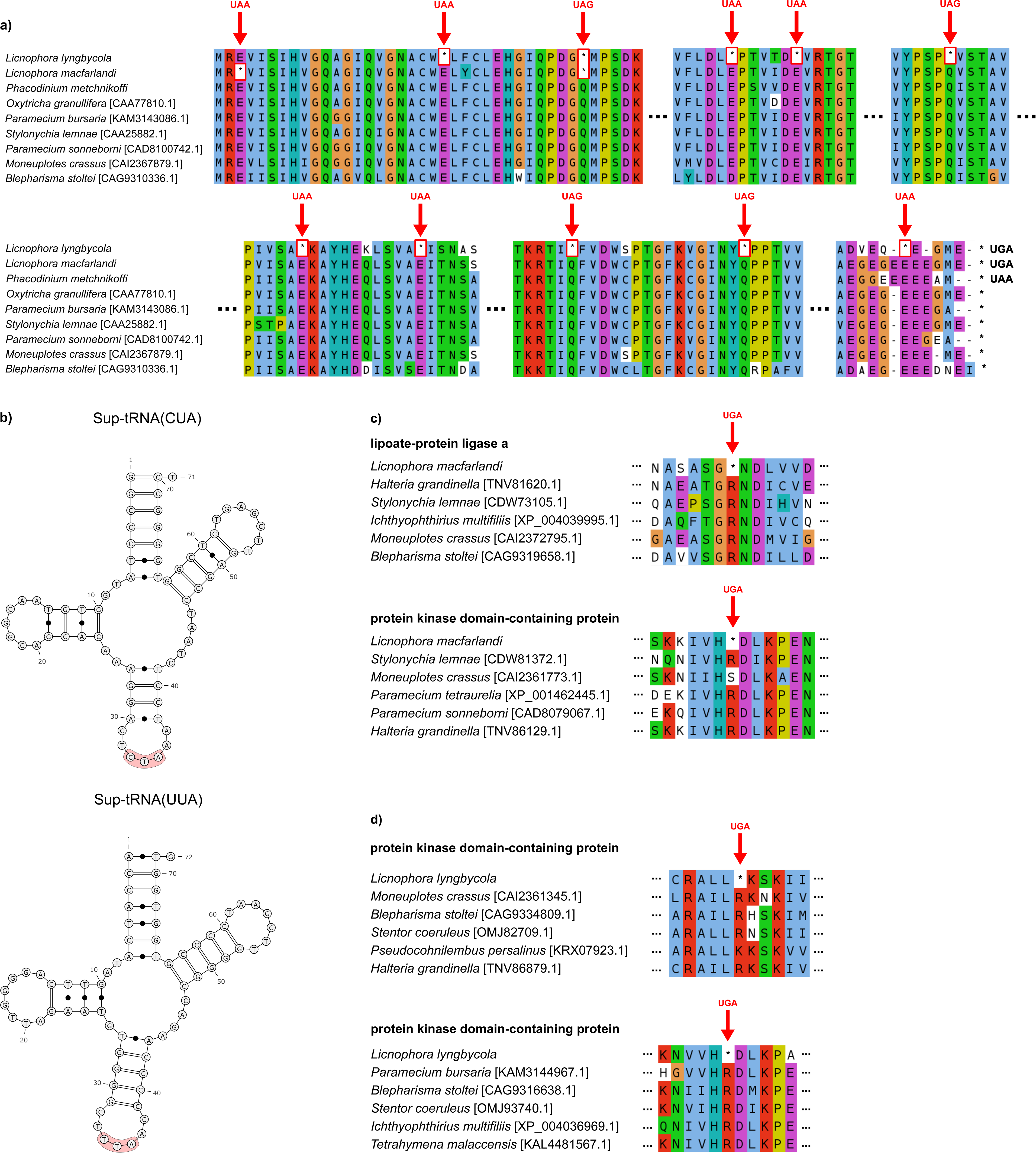
Codon reassignments in *Licnophora* spp. (A) Occurrence of in-frame UAR codons in coding sequences of nuclear genes of *L. macfarlandi* and *L. lyngbycola* exemplified by the gene for α-tubulin; the codon positions are mapped onto a multiple sequence alignment of homologs of both proteins. Both *Licnophora* spp. use exclusively UGA as a stop codon, while *P. metchnikoffi* uses all three standard stop codons (UAA in the example shown). (B) tRNAs cognate to the codons UAG and UAA (see the anticodons CUA and UUA, respectively) identified in the genome assembly of *L. lyngbycola*. The anticodon is highlighted in red. (C) Examples of in-frame UGA codons occurring in coding sequences of nuclear genes of *L. macfarlandi* and *L. lyngbycola*.

Strikingly, Codetta also predicted a reassignment of UGA as an arginine codon in both *Licnophora* spp., but the number of codon positions supporting this altered codon meaning was much lower compared to the respective numbers for the UAG and UAA codons (Table S3). We, therefore, hypothesized that the UGA codon in *Licnophora* spp. may exhibit homonymy, i.e. serving as a sense or stop codon depending on its position within the coding sequence. Dedicated analysis confirmed UGA as a termination codon in *Licnophora* spp. (Fig. S1), but we also identified individual genes where UGA occurs in-frame and most likely specifies arginine (Fig. 1C). Notably, the stop/Arg form of UGA homonymy has been reported from nuclear genomes of various other members of the class Spirotrichea (Chen et al. 2023). The authors did not provide a clear answer regarding the identity of the tRNA decoding UGA as Arg in the spirotricheans concerned. In the supplementary data (their Figure S13) they presented a tRNA with the anticodon UCA identified in the genome of *Uronychia setigera* (a spirotrichean, despite the legend of their figure assigned the species into the class Heterotrichea), but they did not comment on it explicitly in the text. Regardless, our reanalysis revealed that the *U. setigera* tRNA in fact corresponds to the mitogenome-specified tRNA mediating translation of UGA as a tryptophan codon in the mitochondrion (Table S2). Except for an analogous mitochondrial gene for tRNA^Trp^_UCA_ and a putative tRNA_UCA_ pseudogene representing a mammalian contaminant, no candidate nuclear gene for a UGA-cognate tRNA was identified in the *L. lyngbycola* genome assembly either (Table S2). Hence, the mechanism of UGA decoding as Arg in different spirotricheans remains unknown.

### The standard genetic code predominates in the ciliate class Litostomatea

Another segment of the ciliate phylogenetic tree that needs attention when it comes to the genetic code is the class Litostomatea. One lineage often classified as a part of it, the genus *Mesodinium*, was demonstrated early on to exhibit a non-standard genetic code, with UAG and UAA reassigned to tyrosine (Swart et al. 2016; Heaphy et al. 2016). However, phylogenomic analyses have indicated that *Mesodinium* is only distantly related to (other) litostomateans (Lasek-Nesselquist and Johnson 2019; Jiang et al. 2024), supporting a separate class Mesodiniea (Wang et al. 2026). Reports on the genetic code variation in litostomateans (*sensu stricto*) have been contradictory. Single-cell transcriptomes from four phylogenetically diverse litostomateans obtained by Yan et al. (2019) based a conclusion that all three standard termination codons are used normally in these organisms. The standard genetic code has also been claimed for the litostomatean representatives analysed by McGowan et al. (2023), Chen et al. (2023), and Jiang et al. (2024). Likewise, a report on the development of the P10K database ascribed the standard genetic code to most litostomateans included in it, except for unspecified species of the genus *Amphileptus* indicated by the paper to use the NCBI translation table 6 (Gao et al. 2024). We were, therefore, intrigued by two recent studies reporting on genome and/or transcriptome analyses of multiple members of this class, which both concluded that UAG and UGA code for amino acids in all these ciliates (Zhang et al. 2024; Zhou et al. 2024). However, no further details, including the identity of the amino acid(s) putatively encoded by these codons, or what is the molecular mechanism of the changed decoding, were included.

Hence, we set out to clarify the nature of the genetic code in litostomateans by revisiting the genomic and transcriptomic data from this group with a rigorous methodology. We used Codetta to obtain an initial prediction of the genetic code of these organisms, which were further evaluated by manual analyses (Table S1). In most cases Codetta did not indicate any reassignment of any of the three standard termination codons. Furthermore, all the six litostomateans analysed previously by Zhou et al. (2024) and claimed to have reassigned UAG and UGA as sense codons certainly use all three common termination codons as translation termination signals, as revealed by our inspection of samples of transcripts in the respective transcriptome assemblies (Fig. S3). We could not get access to the genome assemblies reported on by Zhang et al. (2024), but the dataset analysed by us does include sequences from nominal close relatives of the respective taxa and they do not exhibit any indication for UAG or UGA being reassigned.

In a few litostomatean genomic datasets UGA was identified as a putative glycine codon by Codetta (Table S1), but we found that these assemblies were contaminated by sequences derived from bacteria of the Absconditabacteria/Gracilibacteria group (Table S4), which is known to have UGA reassigned as a glycine codon (Lukeš et al. 2025). When the bacterial sequences were filtered out, the signal for the UGA reassignment vanished. The transcriptome assembly from *Litonotus* sp. was found to be contaminated by sequences from an *Euplotes* species, explaining the signal for UGA encoding cysteine detected by Codetta. The genome assembly from a *Dileptus* sp. and the transcriptome assembly from an organism denoted *Spathidium* sp. Sr_ci_Sp01 yielded a signal for UAA and both UAR codons, respectively, encoding glutamine (Table S1), but our analysis uncovered that the datasets are contaminated by sequences from other ciliates representing groups known to have these codons reassigned (Table S4). Furthermore, the identification of the latter ciliate as reported in the original publication (Yan et al. 2019) seems to be problematic, as according to our analyses the sequence data do not contain any 18S rRNA sequence attributable to a member of the genus *Spathidium*, yet they contain an 18S rRNA sequence from another litostomatean (most likely attributable to the genus *Lacrymaria*), in addition to the contaminating ciliate (the karyorelictean *Loxodes* sp.; Table S4). Regarding the four genome assemblies of the genus *Amphileptus* included in the P10K database, Codetta did not assign any amino acid to any of the three standard termination codons. Combined, our results do not endorse the previous claims on stop codon reassignments in Litostomatea and the basis of these claims is elusive.

### UGA as an unprecedented leucine codon in the litostomatean Balantidium ctenopharyngodoni

Nevertheless, we did detect a lineage of Litostomatea that has departed from the standard genetic code. It is represented by the species *Balantidium ctenopharyngodoni*, nominally belonging to the polyphyletic genus *Balantidium* but not directly related to the type species *Balantidium entozoon* (Grim and Buonanno 2009; Gürelli and Kesbiç 2025). It seems that the only explicit reflection on the *B. ctenopharyngodoni* genetic code has so far been provided by the authors who generated single-cell transcriptomic data obtained from this species and used it to study its phylogenetic position (Sun et al. 2017). They stated, yet without providing any specific reason for their choice, that they used the “ciliate translation table” (presumably the NCBI translation table 6 with UAG and UAA encoding glutamine), with only UGA considered as a stop codon. More recently, a genome assembly of this species has been reported and analysed in regard to its gene content, but no information on the genetic code employed in the analyses is provided in the paper (Zhao et al. 2024). The results of our analyses of both the transcriptomic and genomic data from *B. ctenopharyngodoni* do indicate an alteration of the genetic code in this ciliate, but that the configuration of codon meanings in this species is exactly reversed compared to the assumption by Sun et al. That is, the data strongly imply that both UAG and UAA serve as termination codons rather than being translated as glutamine (Fig. S4; Table S1). Instead, it is the UGA codon that is reassigned, with a strong signal indicating that the amino acid specified by it is, surprisingly, leucine (Fig. 2A; Table S1).

**Fig. 2.**
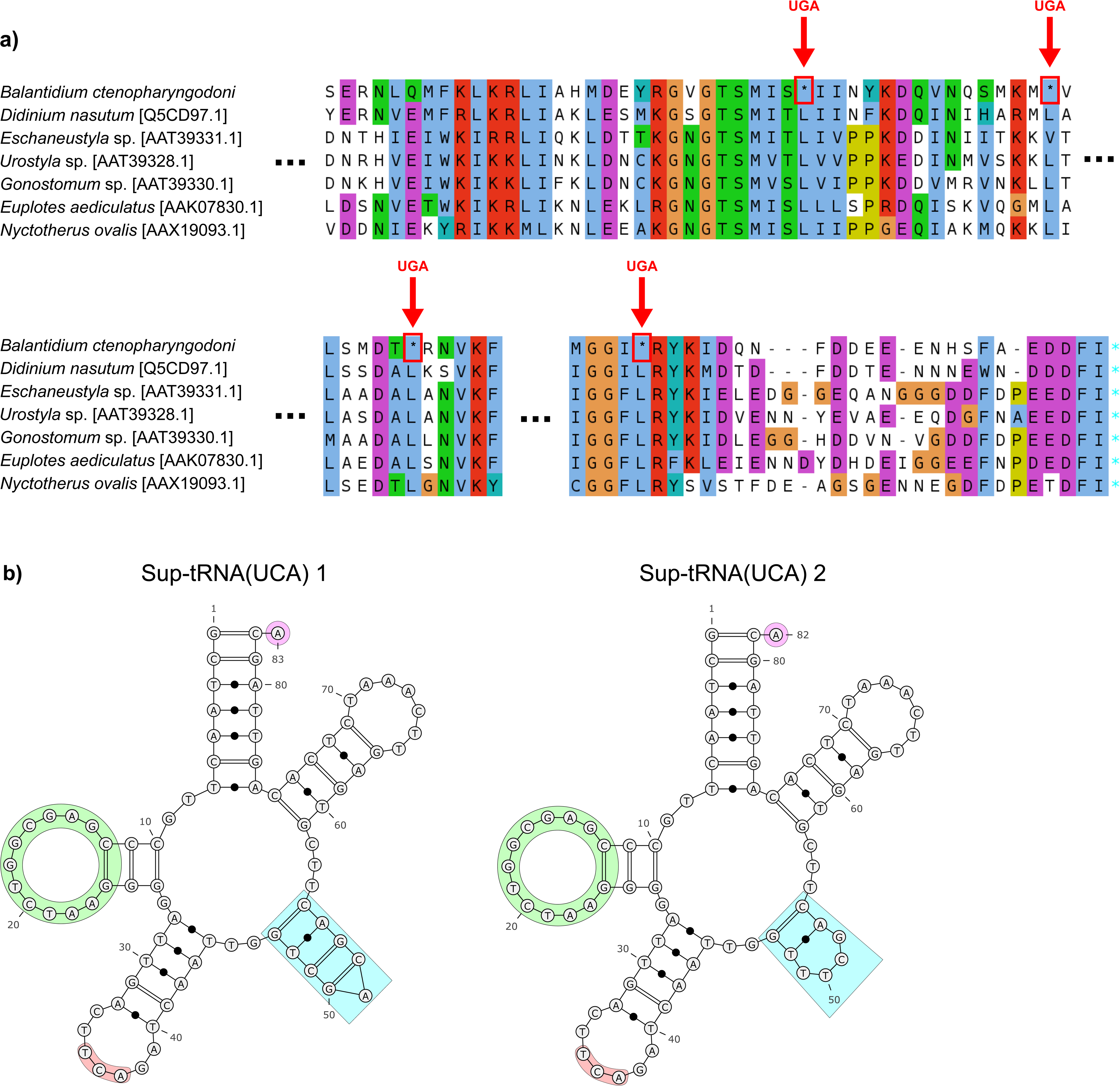
UGA codon reassignment in *Balantidium ctenopharyngodoni*. (A) Occurrence of in-frame UGA codons in coding sequences of nuclear genes of *B. ctenopharyngodoni*, mapped onto a multiple sequence alignment of eRF1. (B) Two different variants of a tRNA specified by the *B. ctenopharyngodoni* genome, exhibiting an anticodon (in red) cognate to UGA and features indicating charging with leucine. The D-loop is identical to the D-loop of a tRNA^Leu^ found in the same genomic data (green) and the variable arm characteristic to tRNA^Leu^ is present (blue). A73, another important tRNA-Leu identity element, is highlighted in pink.

Consistent with our inference, the *B. ctenopharyngodoni* nuclear genome assembly contains two loci specifying slightly different tRNAs with the anticodon UCA cognate to the UGA codon (Fig. 2B). These tRNAs both exhibit features expected in tRNAs charged with leucine (Giegé and Eriani 2023), including a short variable arm, an adenine at the position 73, and a D-loop identical to leucine tRNA also found in the genome (Table S2). Phylogenetic analysis directly supported the emergence of these tRNAs from a standard tRNA^Leu^ ancestor (Fig. S5).

UGA is known to be very flexible in terms of its capacity to specify different translation outcomes: besides translation termination, it serves as a codon underpinning co-translation insertion of selenocysteine (in a homonymous manner, i.e. in a specific sequence context provided by the SECIS element) and has been reassigned to code for various amino acids in “normal” translation events, including tryptophan, cysteine, glycine, serine, and arginine (Lukeš et al. 2025). However, to our knowledge, UGA as a leucine codon has not been described from any translation system so far. To pinpoint the origin of the reassignment in the *B. ctenopharyngodoni* lineage, we investigated its closest relatives. Based on 18S rRNA phylogenies, *B. ctenopharyngodoni* forms a strongly supported clade with *Balantidium polyvacuolum* and a group of marsupial endobionts formally classified as the order Macropodiniida, with the branching order of these taxa unresolved (Gürelli and Kesbiç 2025). There are no genome or transcriptome data available from macropodiniids, but we obtained a transcriptome assembly using RNA-seq reads from *B. polyvacuolum* reported by Bu et al. (2023). The authors did not comment on the genetic code of the organism in their study, and our analysis revealed that *B. polyvacuolum* exhibits the standard genetic code, with UGA serving as a stop codon (Table S1; Fig. S6). We estimated the relative usage of the three synonymous termination codons in *B. polyvacuolum* and found that UGA is employed commonly (∼18% stop codons are UGA), so no evolutionary trend towards its reassignment is apparent in this species. Hence, the stop-to-Leu UGA reassignment observed in *B. ctenopharyngodoni* must have occurred after the divergence of the *B. polyvacuolum* lineage. Whether it might be shared by macropodiniids is a question to be resolved by obtaining genome or transcriptome sequences from any member of the group.

### Updated picture of genetic code evolution in the ciliate superclade SAL

The results presented above correct some of the previous assumptions or claims regarding the genetic code variants present in different lineages of Spirotrichea and Litostomatea. Together with the genus *Mesodinium* (a putative sister lineage of the “core” Litostomatea) and the so-called APM clade comprising several classes of anaerobic ciliates, these classes constitute a broader “superclade” originally known as SAL (Gentekaki et al. 2014), but later also referred to as SAPML (Jiang et al. 2024) or SLAOMP (Wang et al. 2026); here we stick to the original name for simplicity. In Fig. 3 we present an updated scheme of the evolution of the nuclear genetic code in the SAL superclade, reflecting on the previously published robustly documented cases of codon reassignments in this group and the results of our (re)analyses. The genetic code of the anaerobic ciliates constituting the APM clade members has been investigated in depth by Rotterová et al. (2024), who found the standard genetic code as the rule, with a sole known exception represented by *Palmarella salina* (class Armophorea), which has reassigned UAG and UAA as glutamine codons (using only UGA as a termination codon). In addition, Chen et al. (2023) stated that they observed a hundred of UAG codons in coding sequences of *Brachonella spiralis* (another Armophorea representative), without providing the underlying data or proposing the specific meaning of the putative in-frame UAG codons. Our reanalysis indicated that the signal comes from a contaminant, namely sequences derived from a *Blepharisma* sp., so stop-to-Gln UAR reassignment in *P. salina* remains the only known genetic code change in the APM clade. The stop-to-Leu UGA reassignment here reported from *B. ctenopharyngodoni* and the stop-to-Tyr UAR reassignment occurring in the *Mesodinium* lineage are then two additional genetic code alterations in the SAL superclade readily interpretable as lineage-specific switches from the standard genetic code present in the ancestors of these lineages.

**Fig. 3.**
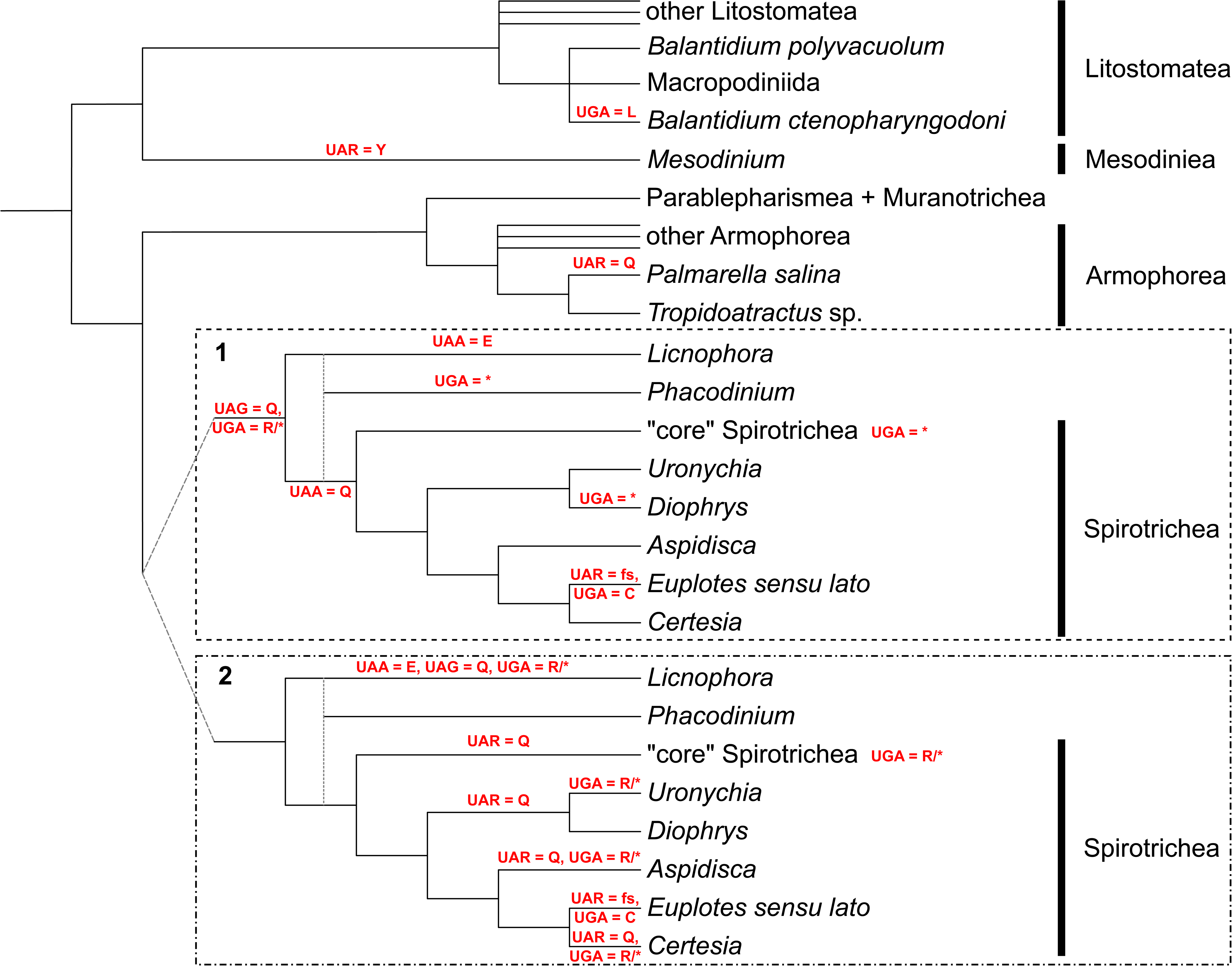
Genetic code changes in the SAL superclade of Ciliophora. The simplified phylogeny is based on a synthesis of multiple molecular phylogenetic and phylogenomic studies (Lasek-Nesselquist and Johnson 2019; Chen et al. 2021; Irwin et al. 2021; Jiang et al. 2024; Rotterová et al. 2024; Gürelli and Kesbiç 2025; Wang et al. 2026). The position of *Phacodinium* relative to other lineages of Spirotrichea varies between analyses, with the two alternative placements (Irwin et al. 2021; Wang et al. 2026) indicated by dashed lines. Inferred changes in the meaning of particular codons are mapped onto branches, following the maximum parsimony principle and additional considerations. For Spirotrichea, a complex history of codon reassignments has to be invoked and no single best scenario can be presently inferred, so two alternatives are presented in the figure: (1) a scenario allowing for reversions of, or secondary changes in, the meaning of particular codons; (2) a scenario with no reversions or secondary changes allowed. The lineage denoted “core” Spirotrichea (encompassing the subclasses Hypotrichia, Choreotrichia, and Oligotrichia) contains representatives that according to the data reported by Chen et al. (2023) vary in the use of the UGA codon, with some of them presumably interpreting it solely as a termination codon while others using it homonymously also as an arginine codon. The labels “UGA=*” and “UGA=R/*” placed to the right of the lineage name in the scenario 1 and 2, respectively, thus indicate recurrent loss of the homonymous decoding and its recurrent establishment (for simplicity, we do not attempt to map these changes within “core” Spirotrichea explicitly). The *Euplotes sensu lato* clade (i.e., Euplotidae) has changed the UAR codons to frameshifting signals (“fs”).

The evolutionary history of the genetic code in the class Spirotrichea is more complex. An early but essentially overlooked study provided evidence suggesting that spirotricheans with UAR codons reassigned to glutamine are not monophyletic to the exclusion of *Euplotes* exhibiting a completely different pattern of codon reassignments (Perez-Romero et al. 2001). This conclusion has been recently corroborated by genome-scale analyses demonstrating that the “euplotid genetic code” (see above) is restricted to the family Euplotidae (*Euplotes sensu lato*; Lobanov et al. 2017; Gaydukova et al. 2023) and this group is phylogenetically nested among spirotricheans with the stop-to-Gln UAR reassignment (Chen et al. 2021). More specifically, the latter genetic code change occurs not only in the phylogenetic core of Spirotrichea comprising the majority of defined orders of the class, but also in non-Euplotidae representatives of the broader order Euplotida that branches as a sister group of the core clade (Fig. 3). Furthermore, as already mentioned above, the homonymy of UGA as a context-dependent arginine-encoding or termination codon has been proposed for some of the spirotricheans with UAG=Gln, notably including the Euplotida member *Uronychia* (Chen et al. 2023). We investigated genome data from multiple Euplotida representatives collectively forming a paraphyletic grade at the base of Euplotidae (Fig. 3; Jiang et al. 2024; Wang et al. 2026) and confirmed the stop-to-Gln UAR reassignment in all of them (Table S1). Furthermore, our analyses also support the notion that UGA is used homonymously to encode arginine or terminate translation in at least two of these lineages, namely the genera *Uronychia* and *Aspidisca*, with a weaker signal of UGA encoding arginine observed also in *Certesia quadrinucleata*, the closest relative of Euplotidae with genome data available whose genetic code has not been previously investigated (Table S3). This finding challenges the previously proposed claim that the genetic code in the more distantly related genus *Diophrys* “may represent an intermediate form between *Euplotes* and *Uronychia*” (Chen et al. 2023), an idea based on the observation by the authors that UGA in *Diophrys* may serve as a low-frequency cysteine codon. In fact, the evidence for this particular UGA meaning in *Diophrys* seems to be weak according to our scrutiny (Table S3), and even this weak signal is likely caused by the data being contaminated by sequences from *Euplotes* (Table S4).

Considering further the results of the genetic code analysis in *Phacodinium* and *Licnophora* presented here, several alternative histories of the genetic code changes during the phylogeny of the class Spirotrichea can be invoked (Fig. 3). In principle, the stop-to-Gln reassignment of the UAG codon and the switch to the stop/Arg homonymy of the UGA codon might have happened in the common ancestor of the whole class, followed by reversion to the standard meaning of both codons in *Phacodinium*. In the common ancestor of Euplotidae, UAG would revert to a termination codon along with spreading inside coding sequences as a frameshifting signal. The UAA reassignment to glutamate in *Licnophora* is an event exclusive to this lineage among spirotricheans, whereas an UAA reassignment to glutamine might have preceded the divergence of Euplotida from the spirotrichean phylogenetic core, with a subsequent further change in UAA meaning in the common ancestor of Euplotidae paralleling the fate of UAG in this lineage. Notably, the scenario assuming a transition of UAG and UAA from glutamine codons to frameshifting signals in *Euplotes* is consistent with observations of UAG or both UAR codons, serving as putative sense codons at a limited number of positions in genes of, respectively, *Euplotes raikovi* (Ricci et al. 2019) and *Euplotes octocarinatus* (Wang et al. 2020). Exactly opposite to the scenario outlined above is refuting the possibility of reversion or secondary reassignment of any codon in the spirotrichean evolution and assuming multiple independent transitions from the standard meaning, including multiple parallel changes inside Euplotida alone. In between these extremes lie histories assuming any combination of reversions and/or secondary reassignments with recurrent primary reassignments. Distinguishing between these possibilities requires a detailed, taxonomically comprehensive evolutionary and functional analysis of the molecular determinants of the varying codon meanings in different spirotrichean lineages, an endeavour that is beyond the scope of the present study.

## Conclusions

Apart from refining the understanding of the genetic code diversity and evolution in ciliates, our results further underscore the significance and surprising commonality of the discrepancies between the actual genetic code variants employed by translation systems of particular taxa and the translation tables assigned to them in databases and genomic studies. It is likely that the literature is replete with artificial results stemming from misinterpretation of nucleotide sequence data in the conceptual translation step. We posit that standardized and rigorous assessment of the codon meaning in the given genome should become a routine part of any studies reporting on new genomic or transcriptomic data.

## Methods

### Compilation of ciliate sequence data and contamination assessment

Transcriptome assemblies from *Licnophora macfarlandi* and *Phacodinium metchnikoffi* generated by Irwin et al. (2021) were downloaded from Dryad (Table S1). Transcriptomic reads for *Spathidium* sp. (SRR10512987), *Balantidium polyvacuolum* (SRR24608340), *Balantidium* cf. *grimi* (SRR27838448), and *Balantidium ctenopharyngodoni* (SRR26318078) were quality-tested with fastqc v0.12.1 (Andrews 2010), then trimmed and assembled with Trinity v2.15.2 (Grabherr et al. 2013) with the -- trimmomatic option. Genomic reads for *Licnophora lyngbycola* (SRR32489309) were quality-tested with fastqc v0.12.1 (Andrews 2010), trimmed with Trimmomatic v0.39 (Bolger et al. 2014), and assembled with SPAdes v3.15.5 (Bankevich et al. 2012).

To assess the taxonomic composition of the genomic and transcriptomic assemblies, SSU rRNA genes were first detected using covariance models representing archaeal, bacterial and eukaryotic SSU rRNAs from the Rfam database v14 (Kalvari et al. 2021) using the cmscan program of the Infernal package v1.1.5 (Nawrocki et al. 2009). These were then annotated by searching against the SILVA database of curated ribosomal RNA sequences v138.1 (Chuvochina et al. 2026) using NCBI BLAST+ blastn v2.17.0 (Camacho et al. 2009). Sequences of interest were then further assessed using manual blastn search against the NCBI nt database. In some cases, however, we observed that the datasets were apparently pre-filtered for some of the contaminating reads, removing prokaryotic 16S reads, but leaving the non-rRNA contaminants.

The decontamination of *Balantidium ctenopharyngodoni* and *B. polyvacuolum* genome assemblies was carried out as follows. Homology searches of all contigs were performed 1) using NCBI BLAST+ blastn v2.17.0 in megablast mode against the NCBI core_nt database (downloaded in April 2025) and 1e-40 e-value cutoff; and 2) using Diamond blastx v2.1.8 (Buchfink et al. 2021) in fast mode against the Uniprot Reference Proteomes database (version 2026_01) using 1e-10 e-value cutoff. For each contig the best BLAST hit was established based on the blast score. Blastn hit scores were divided by three to allow direct comparison with the blastx hits. Contigs with the best hit outside of ciliates with >= 80% sequence identity were removed and contigs without significant hits with GC content outside the 22% - 41% range were also removed. The enrichment of ciliate contigs of the litostomatean lineages with a suspected Absconditabacteria/Gracilibacteria prokaryotic contamination was also carried out using homology searches as outlined above, but with different selection criteria, specifically removing all contigs without any significant BLAST hit or with the best BLAST hit outside of Ciliophora.

### Analyses of the genetic code and codon usage

Automated inference of the genetic code of all organisms mentioned was performed using Codetta v2.0 (Shulgina and Eddy 2023) and, in some cases, FACIL (Dutilh et al. 2011). To infer the relative usage of the UGA stop codon in *Balantidium polyvacuolum*, we first used TransDecoder2 v1.0.8 (Mao et al. 2025) to define coding sequences in the assembled transcript contigs. We included the optional second step using homology searches as ORF retention criteria. Specifically, using the output of TD2.LongOrfs, we searched the Swissprot database (downloaded 26/2/2026) with diamond blastp v2.1.8, and the Pfam-A (downloaded 2/5/2020) database using HMMER hmmscan v3.4 (Eddy 2011). Outputs of these searches were then used as an input for the TD2.Predict step. Termination codons in coding sequences were counted from the TD2.Predict output using a custom script.

### Analyses of tRNA genes

Identification of tRNA genes in all relevant genome assemblies was performed using tRNAscan-SE v2.0.12 (Chan et al. 2021) with the -E option. For presentation purposes, the secondary structures predicted for tRNAs of special interest were visualized using VARNA v3.93 (Darty et al. 2009) and modified in Inkscape v1.2.2 (www.inkscape.org). All tRNA sequences found in the *B. ctenopharyngodoni* genome were aligned with MAFFT v7.505 (Katoh and Standley 2013), and the alignment was inspected for any irregularities. Based on the inspection, tRNA^Pro^_AGG_ from contigs_11633:728-819 (-) was removed, because it had suspicious insertions in its sequence. The sequences were aligned again, and the automatically generated alignment of tRNA^SeC^_TCA_ sequences was manually adjusted to correctly align to the anticodon region. Finally, the anticodon was removed from the alignment. IQtree v2.4.0 (Minh et al. 2020) was used for building the phylogenetic tree, employing the K2P+G4 model determined by ModelFinder (Kalyaanamoorthy et al. 2017), with 100 standard nonparametric bootstrap replicates.

For the evolutionary analysis of tRNA genes in Spirotrichea, we chose *Euplotes aediculatus* (GCA_984074515.1), *Stylonychia lemnae* (GCA_000325865.3), *Oxytricha trifallax* (O_trifallax_2020 from the OxyDB (https://oxy.ciliate.org/downloads.php)), *Certesia quadrinucleata*, *Aspidisca* sp., *Diophrys appendiculata*, *Uronychia setigera*, *Phacodinium* sp. (GWHDPZM00000000.1), and *Licnophora lyngbycola*; each genome was searched for eukaryotic tRNAs using tRNAscan-SE (see above). The results were pooled together, removing all suspected pseudogenes (except for NODE_96585_length_243_cov_0.873494.trna1 from Licnophora lyngbycola, as it was predicted to be a suppressor tRNA) and tRNAs that did not have an amino acid determined. Next, the sequences were aligned with MAFFT v7.505 (Katoh and Standley 2013), and manually inspected for artefact sequences and introns. Two sequences were removed (NODE_16214_length_868_cov_1.085967.trna1 from *L. lyngbycola* and JAGYYM010005468.1.trna2 from *C. quadrinucleata*). All obvious introns were also removed from the sequences, based on online tRNAscan-SE annotation. The whole dataset was realigned using the same parameters. From this alignment, well-aligning sequences were chosen as seed and the dataset was realigned again with the --seed option. Anticodons were manually deleted from the alignment, and a phylogenetic tree was built using IQtree v2.4.0 (Minh et al. 2020), using the GTR+F+R7 model found by ModelFinder (Kalyaanamoorthy et al. 2017), with 1000 replicates of the SH-like approximate likelihood ratio test (Guindon et al. 2010) and 1000 ultrafast bootstrap replicates (Hoang et al. 2018).

## Supplementary Material

**Fig. S1.** Multiple sequence alignments of chosen proteins showing that *Phacodinium metchnikoffi*, although rarely, uses UGA as a termination codon.

**Fig. S2.** Phylogenetic analysis of tRNA genes in *Liconophora lyngbycola* and other selected spirotrichean species.

**Fig. S3.** Multiple sequence alignments of selected proteins showing that the litostomateans with the standard genetic code indeed use all three stop codons for translation termination.

**Fig. S4.** Multiple sequence alignments of selected proteins showing that *Balantidium ctenopharyngodoni* uses both UAA and UAG as termination codons.

**Fig. S5.** ML phylogeny of tRNAs found in the *Balantidium ctenopharyngodoni* genome.

**Fig. S6.** Multiple sequence alignments of chosen proteins showing that *Balantidium polyvacuolum* uses the UGA codon for translation termination.

**Table S1.** Ciliate species analyzed in this study and their respective genetic codes predicted by Codetta.

**Table S2.** A list of relevant tRNAs found in the genomes of *Balantidium ctenopharyngodoni* and *Licnophora lyngbycola*, along with tRNAs reported in Chen et al. (2023).

**Table S3.** Codetta log decoding probabilities from selected organisms in our study.

**Table S4.** A list of selected sequences of SSU rRNA genes and protein-coding genes representing target ciliates and contaminating taxa.

## Acknowledgements

We thank Jana Salamonová for some of the initial genetic code analyses in ciliates.

## Author contributions

E.M. performed a major part of the bioinformatic and phylogenetic analyses, prepared figures and some supplementary items, and contributed to the writing. V.Ž. discovered the UGA reassignment in *B. ctenopharyngodoni*, carried out part of the bioinformatic analyses, and prepared some of the supplementary items. M.E. conceived the research, secured funding, supervised E.M., contributed to bioinformatic analyses, and drafted most of the manuscript.

