## Supplementary figures for "An unprecedented codon reassignment and two new genetic code variants in ciliates"

**This file contains supplementary Figures S1 to S6.**

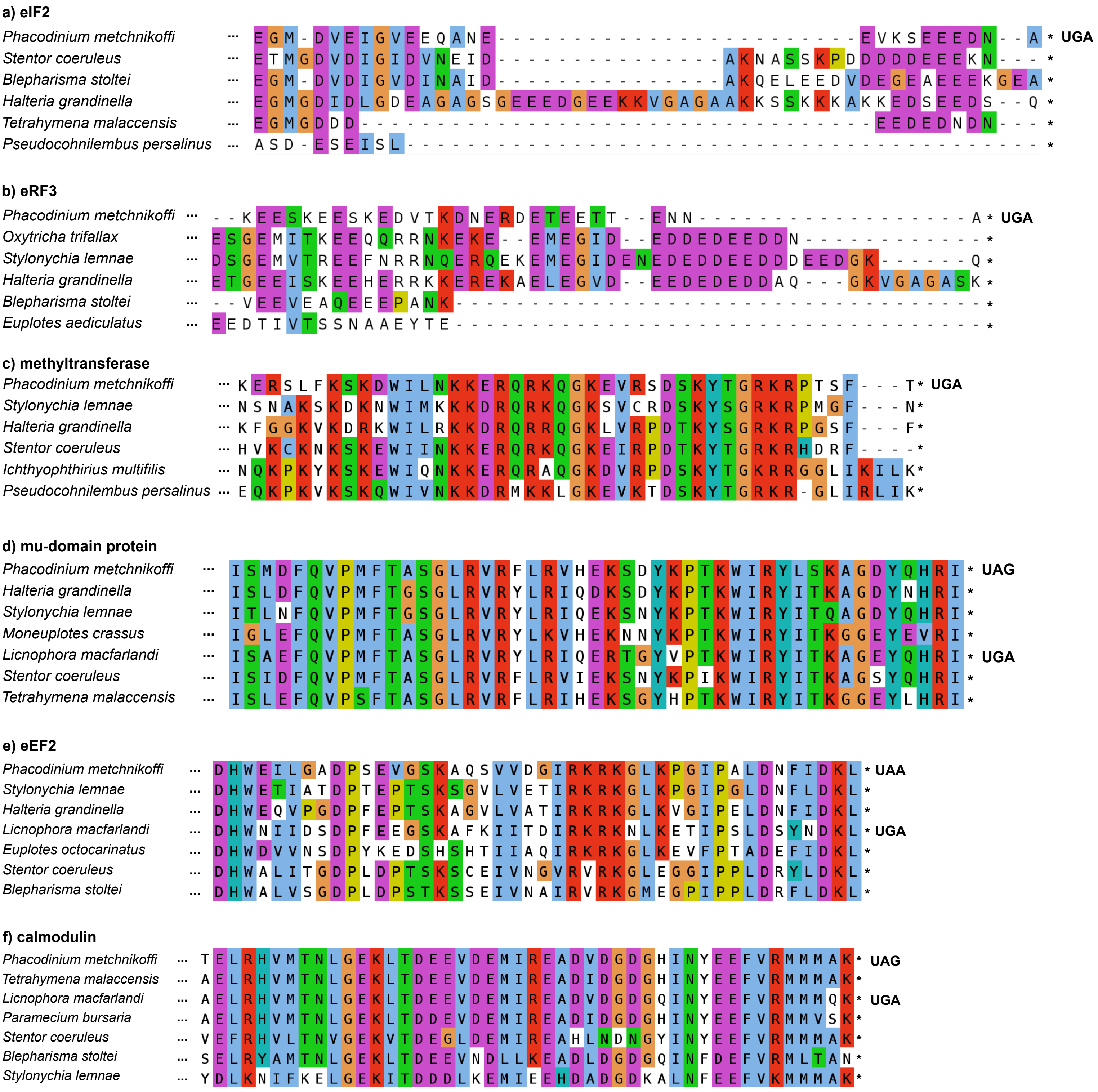


Fig. S1: Multiple sequence alignments of chosen proteins showing that *Phacodinium metchnikoffi*, although rarely, uses UGA as a termination codon. On the other hand, UGA is the sole termination codon in *Licnophora macfarlandi*.

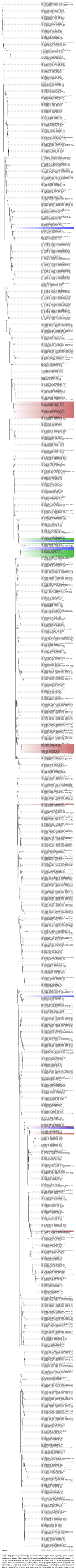


Fig. S2: Phylogenetic analysis of tRNA genes in *Liconophora lyngbycola* and other selected spirotrichean species. Sequences used as seed for the final alignment (see Methods) are marked as "_seed_". Branches with SH-aLRT support and ultrafast bootstrap values being >80 and >95, respectively, are marked with a periwinkle circle. tRNAs with anticodons of interest (UUA, CUA and UCA) are highlighted in blue, green, and red, respectively. As expected, $tRNA^{SeC}_{UCA}$ sequences clustered together, except for one of the *L. lyngbycola* SeC tRNAs, which is likely a bacterial contaminant. Likewise, the sequences of $tRNA^{Sup}_{UCA}$, which decodes UGA as tryptophan in ciliate mitochondria, cluster together. The $tRNA^{Sup}_{UCA}$ from *Euplotes aediculatus* that reads UGA as Cys clusters together with other Cys tRNAs. The two remaining $tRNAs^{Sup}_{UCA}$ are possibly contaminants. Concerning all $tRNAs^{Sup}_{CUA}$ sequences, the situation is pretty straightforward; they cluster together with one another in a $tRNA^{Gln}$ group, suggesting that they were already present in the common ancestor of all represented species. However, the origins of $tRNAs^{Sup}_{UUA}$ genes are more difficult to pinpoint. In *Uronychia setigera*, *Stylonychia lemnae* and *Oxytricha trifallax*, these tRNAs likely have the same origin as $tRNAs^{Sup}_{CUA}$. The $tRNA^{Sup}_{UUA}$ from *Aspidisca* sp. clusters together with $tRNA^{Leu}$ sequences; however, Codetta results are quite definitive about the fact that UAA is decoded as glutamine in this organism. There is also an odd $tRNA^{Sup}_{UUA}$ in *E. aediculatus*, which is part of a clade of tRNAs with varying anticodons and predicted amino-acid identities. The online version of tRNAscan-SE assigned the highest score to Arg, with Trp and Cys as close contenders. The *L. lyngbycola* $tRNA^{Sup}_{UUA}$ presumably employed to decode UAA as glutamate branches separately, but not too far from other glutamate tRNAs.

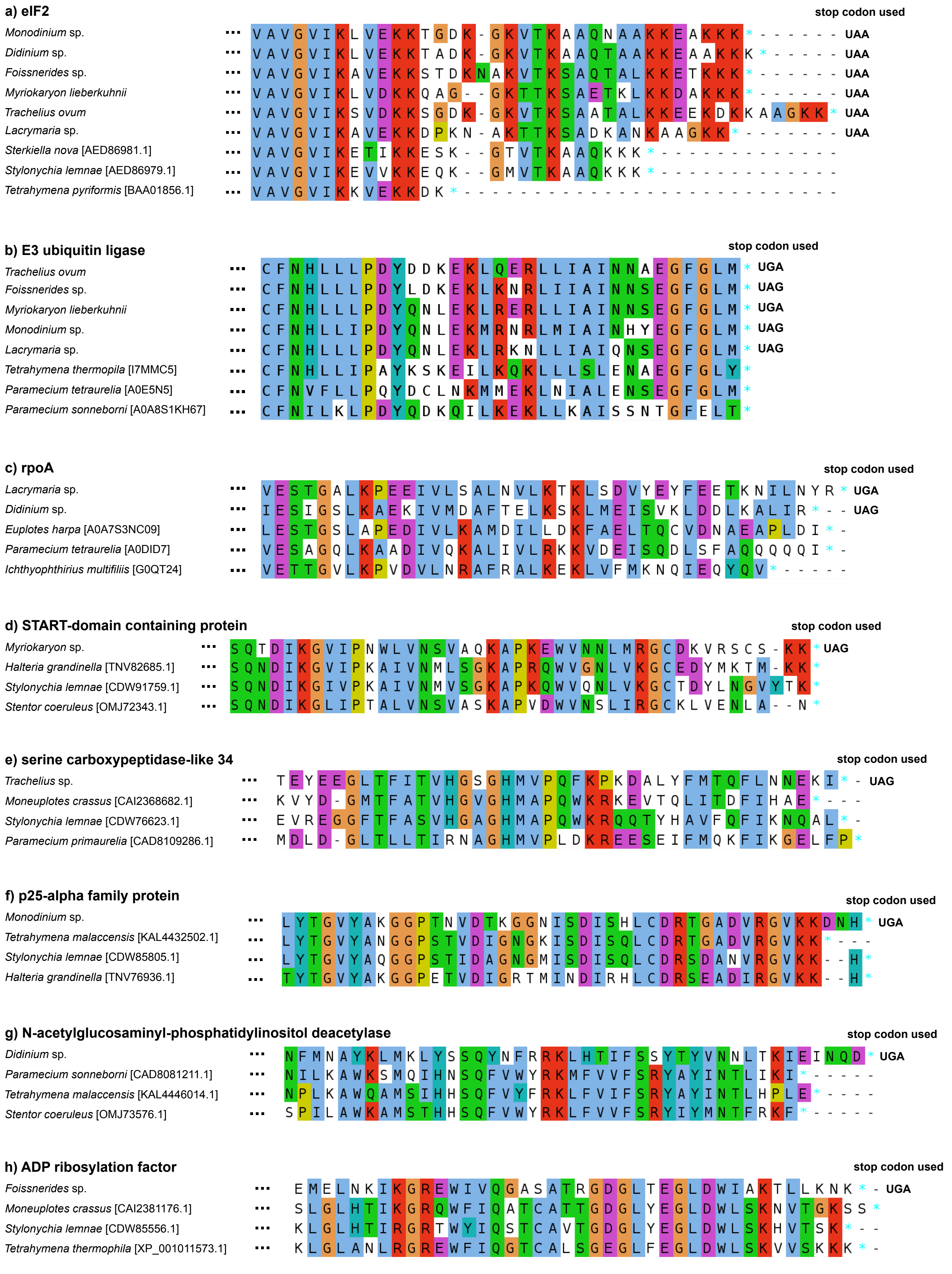


Fig. S3: Multiple sequence alignments of selected proteins showing that the litostomateans with the standard genetic code indeed use all three stop codons for translation termination.

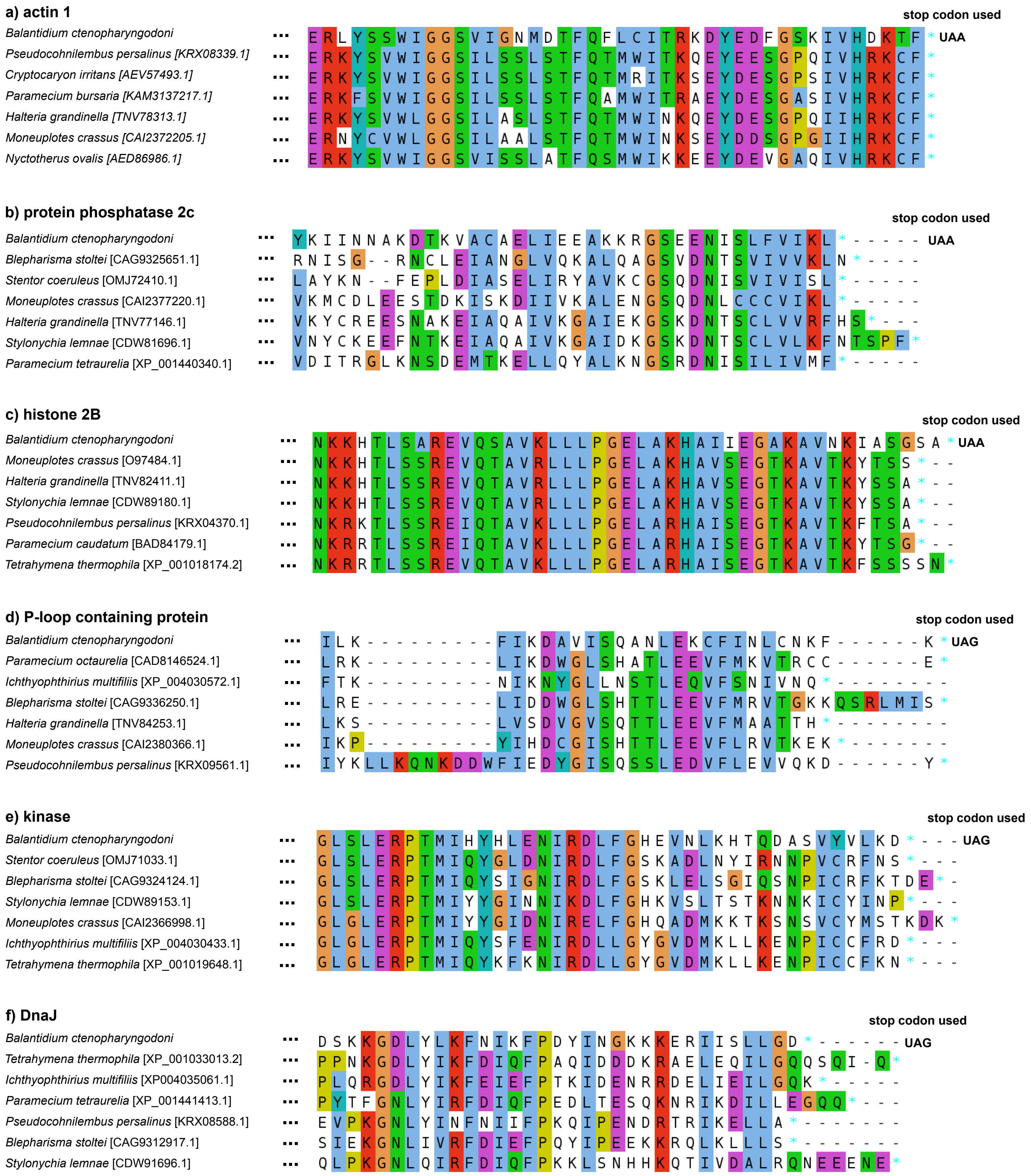


Fig. S4: Multiple sequence alignments of selected proteins showing that *Balantidium ctenopharyngodoni* uses both UAA and UAG as termination codons.

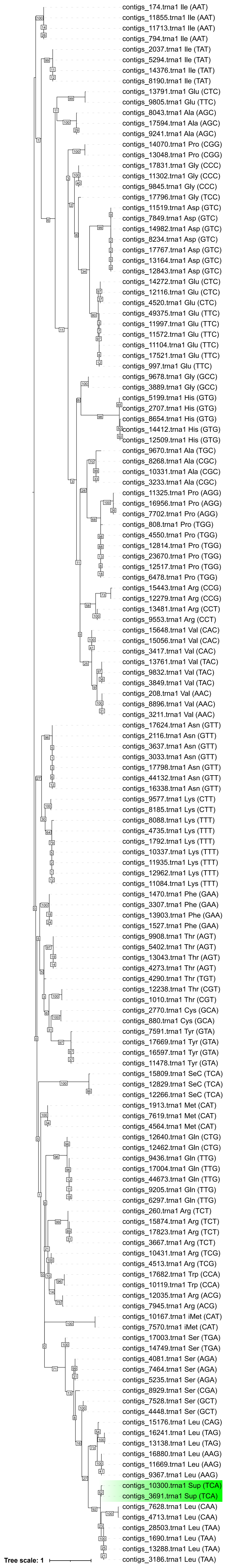


Fig. S5: ML phylogeny of tRNAs found in the *Balantidium ctenopharyngodoni* genome. tRNAs which were predicted as suppressor with UCA anticodon form a clade within leucine tRNAs (highlighted in green).

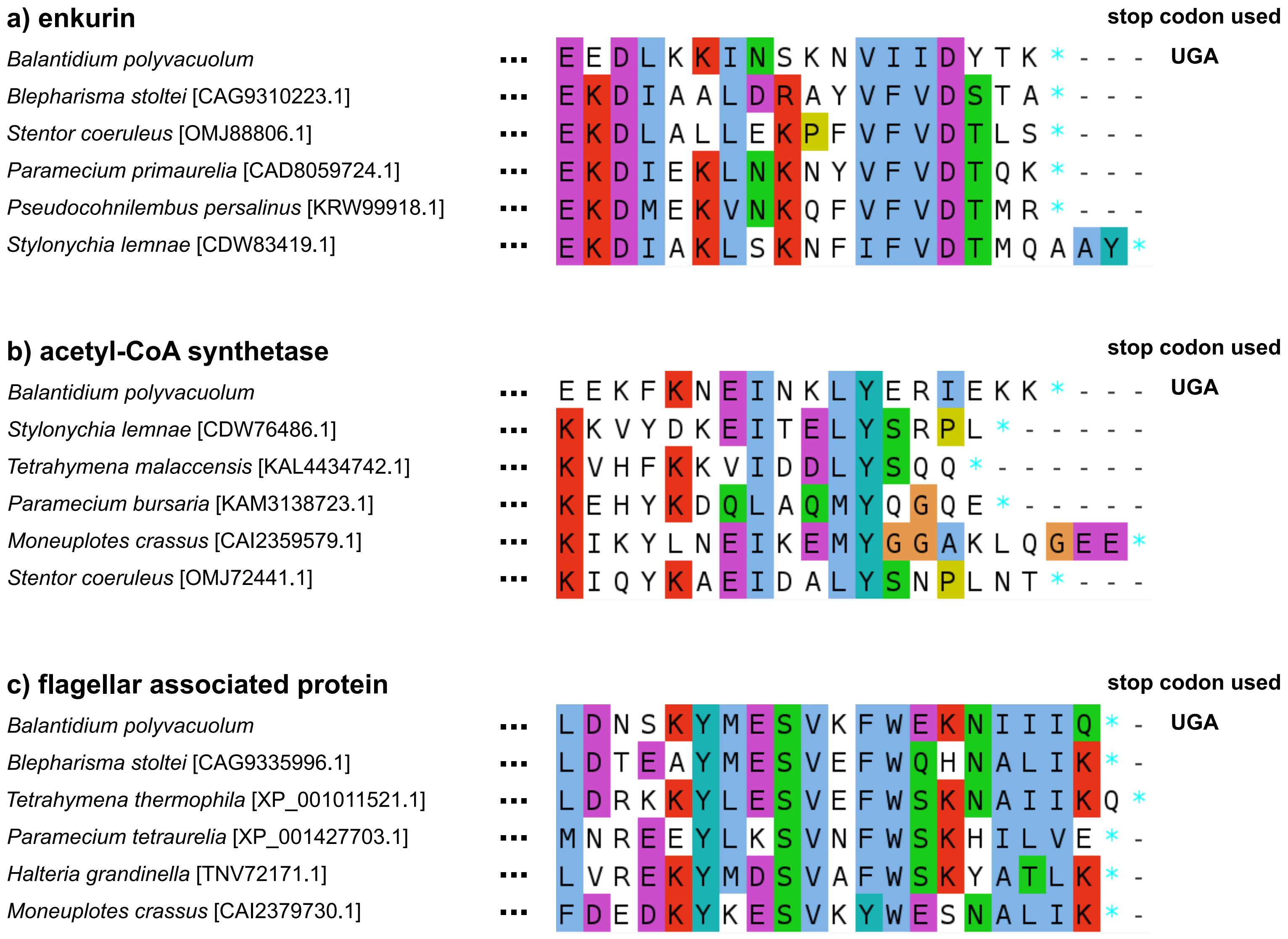


Fig. S6: Multiple sequence alignments of chosen proteins showing that *Balantidium polyvacuolum* uses the UGA codon for translation termination.
